# Cholecalciferol improves cognitive impairment by amending impaired insulin signaling in sporadic Alzheimer’s Disease Rats

**DOI:** 10.1101/2023.02.27.530250

**Authors:** Tushar Kanti Das, Estinnorell Yong, Mas R.W. Abdul Hamid

## Abstract

Alzheimer’s disease (AD) is the most common form of dementia and contributes to 50-70% of neurodegenerative brain diseases. AD has been associated with poor vitamin D nutrition, which is correlated with low mood and impaired cognitive performance in older people. The impact of vitamin D on the insulin signaling pathway in AD is not well known. Hence, this study was to explore the effects of cholecalciferol (Vitamin D_3_) on the expression of IRS-1, IRS-2, Akt, pAkt (Ser473), and GLUT3 in the sporadic AD rat model. The rats were induced to develop sporadic AD by intraperitoneal administration of Scopolamine. The downregulation expression of IRS-1, IRS-2, Akt, pAkt (Ser473) and GLUT3 may lead to impaired insulin signaling which is associated with the development of AD. All these data were compared to Saline-treated control rats. However, cholecalciferol treatment in AD rats may improve memory performance by increasing the expression of insulin signaling proteins and hence ameliorates impaired insulin signaling. All these data were compared to Scopolamine–induced AD rats and sunflower oil-treated rats. Therefore, cholecalciferol treatment may be an alternative approach for the treatment of AD.

## Introduction

Alzheimer’s disease (AD) is the most common form of dementia in elderly individuals and is associated with progressive neurodegeneration of the human neocortex. At the present time, AD is a foremost global health problem in worldwide. Around 6.2 million Americans are diagnosed with AD and it will be increased abound 13.8 million in 2060. During covid-19 pandemic death rate with AD is increased by 16% in the USA [1]. Although more than 100 years ago AD was first identified, the precise biological changes of AD, and how it can be prevented or alleviated are still mostly mysterious [2]. Therefore, studying the mechanism of AD pathogenesis and finding a new treatment strategy for the prevention and/or remedy of AD are one of the most relevant challenges in AD research.

Patients with AD have a high prevalence of vitamin D deficiency, which is also associated with low mood and impaired cognitive performance in older people. Numerous epidemiologic and genetic studies indicate that vitamin D deficit can cause cognitive impairment such as impaired memory and learning [3,4] with hypovitaminosis D [5,6], vitamin D receptor (VDR) polymorphisms and dysregulated VDR mRNA [7,8]. Previous studies documented that prenatal vitamin D deficiency disrupts brain development and alters the expression of growth factors and neurotrophin receptors in the adult dentate gyrus [9]; maternal vitamin D deficiency alters neurogenesis in developing brains [10-12] adult hypovitaminosis D increases the proliferation of neuroblasts in the sub-granular zone of the hippocampus and alters their neuronal differentiation [13,14]. Therefore, Vitamin D is suspected to be a potential modulator of neurogenesis.

The association between vitamin D status and glucose intolerance has been known since the early 1990s [15-17]. It has been shown that the prevalence of hypovitaminosis D is higher in diabetic patients than in non-diabetic people [18]. Indeed, vitamin D improves insulin resistance by regulating insulin sensitivity, β-cell function [19] and gene transcription associated with insulin-growth factor-1 (IGF-1) [20]. Converging evidence proposed insulin signaling impairment association with the pathogenesis of AD. Recent study suggested that defective glucose transportation is associated with insulin signaling in AD brain [21, 22]. Despite these studies, nothing is known about the impact of vitamin D on the insulin signaling pathway during aging or in neurodegenerative disorders such as AD. To further elucidate the potential roles of Vitamin D_3_ (cholecalciferol), we studied the effect of cholecalciferol on the scopolamine-induced AD rat model.

Scopolamine acts as a competitive antagonist for acetylcholine muscarinic receptors. It binds to acetylcholine receptors disrupting acetylcholine neurotransmission and regulation of cognitive function which can lead to memory impairment in animals. It is commonly used for the study of cognitive deficiency in animal models of AD [21-24]. Therefore, in this study, we investigated the effects of cholecalciferol on the mediators of insulin signaling proteins such as IRS-1, IRS-2, Akt,, p-Akt (Ser473); glucose transporter GLUT3 in a rat model of sporadic AD which was generated by intraperitoneal (i.p.) injection of Scopolamine.

## Materials and Methods

### Animals

24 male Sprague Dawley rats aged four months old and weighed 350 ± 50 g provided by PAPRSB Institute of Health Sciences, Universiti Brunei Darussalam. The rats were provided standardized food (Specialty Feeds, Western Australia) and water and, and were maintained under the standard laboratory conditions (12h light/dark cycle). The animal handling and experiments were performed in accordance with the institutional guidelines (UBD/AVC-R1/1.26) by PAPRSB Institute of Health Sciences Research Ethics Committee and Universiti Brunei Darussalam Research Ethics Committee.

### Materials

Scopolamine and cholecalciferol were purchased from Sigma-Aldrich, USA. Sunflower oil was purchased from a retail store in Brunei Darussalam. Immunohistochemistry kit (R.T.U. Vectastain Universal Elite ABC Kit and DAB kit) was purchased from Vector Laboratories (CA, USA). Pierce™ BCA Protein Assay Kit and ECL advanced reagent kit were purchased from Thermo Fisher Scientific (MA, USA) and Bio-Rad (CA, USA), respectively. Protease inhibitor tablets were purchased from Roche Applied Science (Mannheim, Germany). Antibodies against IRS-1, IRS-2, Akt (pan) and p-Akt (Ser 473) were purchased from Cell Signaling Technology (Danvers, MA, USA). Antibodies against β-Actin was purchased from Upstate Biotechnology (Lake Placid, NY, USA) and Sigma Aldrich (St. Louis, MO, USA) respectively. All analytical reagents were purchased from Merck, Germany and Sigma-Aldrich, USA.

### Drug interventional studies

The rats were randomly divided into four groups as follows; (a) control (saline-treated), (b) scopolamine-induced (c) oil-treated (sunflower oil as vehicle) and (d) scopolamine+cholecalciferol (Table 1). Control group was administered with 0.9% saline (w/v) and scopolamine (2.5mg/kg) was administered to all the groups through i.p route for 28 consecutive days. The treatment groups were given cholecalciferol (71.40 IU/kg) by oral gavage for 28 consecutive days. Cholecalciferol was dissolved in sunflower oil. Sunflower oil was administered orally via oral gavage. Our study did not observe any signs of suffering in animal, although oral gavage was an invasive procedure.

**Table 1.**
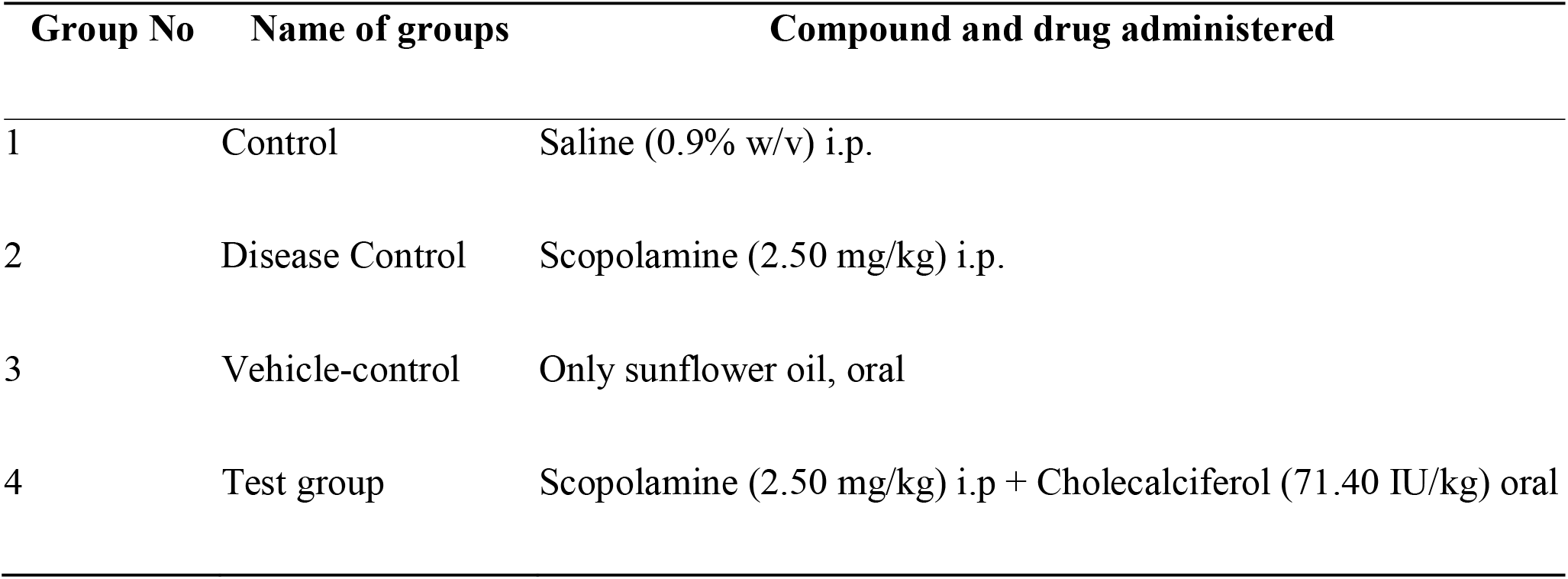
Experimental design and drug doses in different groups of rats.

| Group No | Name of groups | Compound and drug administered |
| --- | --- | --- |
| 1 | Control | Saline (0.9% w/v) i.p. |
| 2 | Disease Control | Scopolamine (2.50 mg/kg) i.p. |
| 3 | Vehicle-control | Only sunflower oil, oral |
| 4 | Test group | Scopolamine (2.50 mg/kg) i.p + Cholecalciferol (71.40 IU/kg) oral |

### Rectangular maze test

The maze consisted of a completely enclosed rectangular box with an entry and reward chamber attached at the opposite ends. The box was partitioned with wooden slats into blind passages leaving just twisting corridor leading from the entry to the reward chamber. Transfer latency (the time taken to reach the reward chamber) was recorded beginning from day 1 till day 28. Three readings were taken per animal, mean was calculated and indicated as learning score (transfer latency) for that animal. Lower scores indicated efficient learning while higher scores indicated poor learning in animals

### Collection of brain tissue

After 28 days, the rats were anesthetized with an intraperitoneal (i.p.) injection of Avertin (300 mg/kg) and transcardially perfused with PBS followed by 4 % paraformaldehyde. The brains were collected and washed in chilled 1x PBS (pH 7.4) (Bio-Rad, CA, USA) for three times. Each group consisting of six brains (Table 1) where three brains were used for immunohistochemistry (IHC) and the remaining three brains were used for immunoblotting from each group of rats.

### Immunohistochemistry (IHC) and scoring

5µm thickness brain tissue sections from different groups of rats (*n* = 3 at each time point for each rat) were used for IHC using R.T.U Vectastain Universal Elite® ABC Kit (Vector Laboratories, USA) using standard protocol. Tissue sections were incubated with primary antibodies Akt and p-Akt (Ser473) at 4°C overnight. Subsequent procedures were performed according to the manufacturer’s instruction (Vector Universal elite® ABC kits, Vector Laboratories, USA). Finally, tissues were counterstained with Hematoxylin, dehydrated and mounted. These sections were observed using a light microscope with lens objective 40x (Olympus, USA). A series of four representative images per section was collected. IHC analyses were scored according to the intensity of staining and percentage of positive cells stained with respective antibodies. The scoring was done as follows: 0, <5%(negative); 1,5–25%(weak); 2, 25–50% (moderate); 3, 50–75% (strong), and 4, >75% (very strong) [23]. The analysis (*n* = 3) was done in triplicate.

### Preparation of brain lysates

The remaining three brains from each group were used for immunoblotting. Each brain with known weight was homogenized at 4°C using automated homogenizer and with IX RIPA buffer containing protease inhibitor tablet (Roche, Germany). After that the brain lysates were centrifuged at 12,000 rpm for 10 mins, and 4°C to obtain the supernatants which were stored frozen in -80°C for further studies. Protein concentrations were determined using Pierce™ BCA Protein Assay Kit (Thermo Fisher Scientific).

### Immunoblotting

50µg of total brain lysate was boiled at 95°C for 10 min in 4X gel-loading buffer (Novex, USA). The protein extract was separated by gel electrophoresis, and transferred onto nitrocellulose membrane (Amersham Bioscience, USA), blocked, and probed with their respective primary antibodies. The protein bands were visualized by chemiluminescence using Clarity™ Western ECL Substrate (Bio-Rad). Blots were digitally developed using VersaDoc™ imaging system (Bio-Rad). Pre-stained Precision Plus™protein standards (Bio-Rad) were used to estimate the apparent molecular weight of the protein bands. β-actin (Sigma, USA) was used as an internal control in all analyses to ensure comparable protein loading. The phospho-proteins levels were normalized with corresponding total protein. Densitometry analysis was performed by using NIH ImageJ software to measure the optical densities of the targeted protein band and normalized to the intensity of the endogenous β-actin level.

### Statistical analysis

Data are expressed as mean ± standard deviation (SD). Data were analysed using ANOVA followed by post-hoc test using SPSS 20.0 (SPSS Inc., Chicago, Illinois, USA). Statistical significance was set at p<0.05.

## Results

### Abridged spatial memory in scopolamine-induced AD rats and noticeable reversal by cholecalciferol

The spatial memory test was performed using the rectangular maze (Figure 1). The transfer latency was significantly reduced (*p*<0.001) in saline treated rats, demonstrating higher efficiency of learning in saline-treated rats. However, transfer latency was significantly increased (*p*<0.001) in scopolamine-induced AD rats during this time period, indicating progressive loss of efficiency of learning in scopolamine-induced AD rats. Additionally, transfer latency was progressively decreased (*p*<0.001) in cholecalciferol-treated AD rats during this time period (Figure 1).

**Figure 1.**
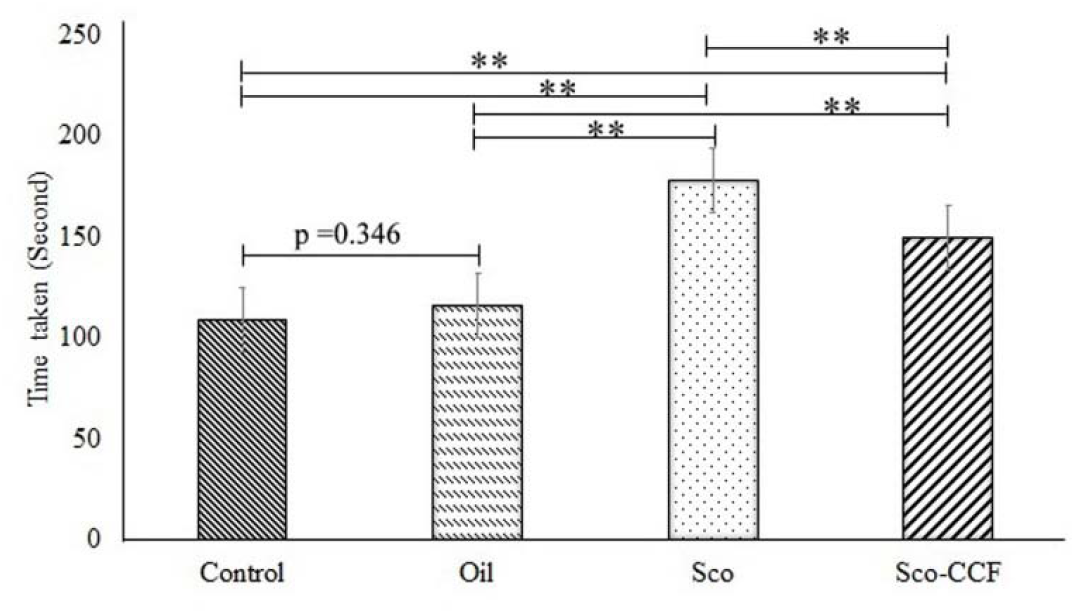
Time taken for rectangular maze test by four groups of rats (n=6). Memory impairment was induced by scopolamine however improved by cholecalciferol in AD rats. Data expressed as mean ± SD. Control: saline-treated, Oil: Sunflower oil treated, Sco: Scopolamine and Sco-CCF: Scopolamine-cholecalciferol treated. *P < 0.01 and **P < 0.001.

### Reduced number of immunostained cells positive for hallmark proteins in scopolamine-induced AD rat brain and marked reversal by cholecalciferol

Immunohistochemistry was used to detect the protein expression of Akt and p-Akt (Ser 473) in the cortex region of rat brains (Figure 2). From the IHC scoring analysis, it was found that the expression of Akt and p-Akt (Ser 473) were significantly decreased in AD rats compared to saline-treated rats. Treatment with cholecalciferol in AD rats significantly increased the levels of these proteins.

**Figure 2.**
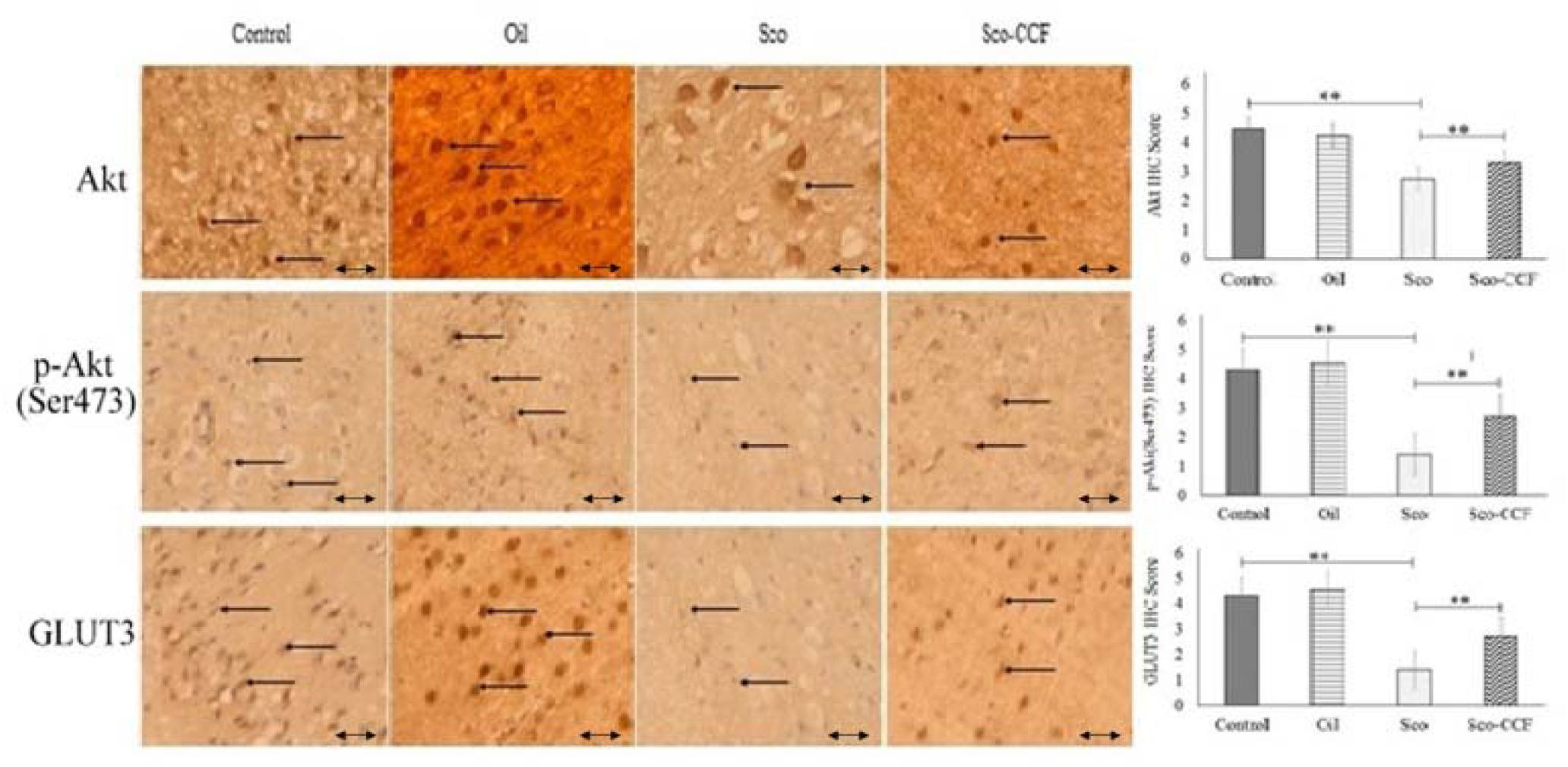
Immunohistochemistry images of (i) Akt (ii) p-Akt (Ser473) and (iii) GLUT3 in the brain sections (cortex) of rat that were treated with Control: saline-treated, Oil: Sunflower oil treated, Sco: Scopolamine and Sco-CCF: Scopolamine-cholecalciferol. The assay (n = 3) was done in triplicate. Scale bar represents 1 μ. Lens objective: 40x. (**p < 0.01).

### Abridged levels of upstream insulin signaling proteins in AD rat brain and cholecalciferol significantly upturned it

To examine the expression of major upstream insulin signaling proteins in the brain involved, we investigated the protein levels of IRS-1, IRS-2, Akt and p-Akt (Ser 473) in AD-rat brain homogenates by immunoblotting (Figure 3). Protein expression of IRS-1 and IRS-2 in scopolamine-induced group, were significantly reduced (P<0.001) compared to saline-treated group. Cholecalciferol did not significantly increase the expression of both IRS-1 (P=0.497) and IRS-2 (P=0.611) in scopolamine-induced group (Figure 3a). In addition, it shows significant difference (P<0.001) in the expression of IRS-1 between oil-treated and saline-treated group (Figure 3a) and the expression of IRS-2 did not differ significantly (P=0.560) between oil-treated and saline-treated group. Expression of Akt (Figure 3c) and pAkt (Ser473) (Figure 2 & 3d) proteins in scopolamine-induced group, were significantly reduced (P<0.001), compared to saline-treated group. However, cholecalciferol significantly increases (P<0.001) the expression of both Akt and pAkt (Ser473) in scopolamine-induced group. In addition, Akt expression was significantly increased (P<0.001) and pAkt (Ser473) did not differ significantly (P=0.184) between saline-treated group and sunflower oil-treated rat brain (Figure 3d).

**Figure 3.**
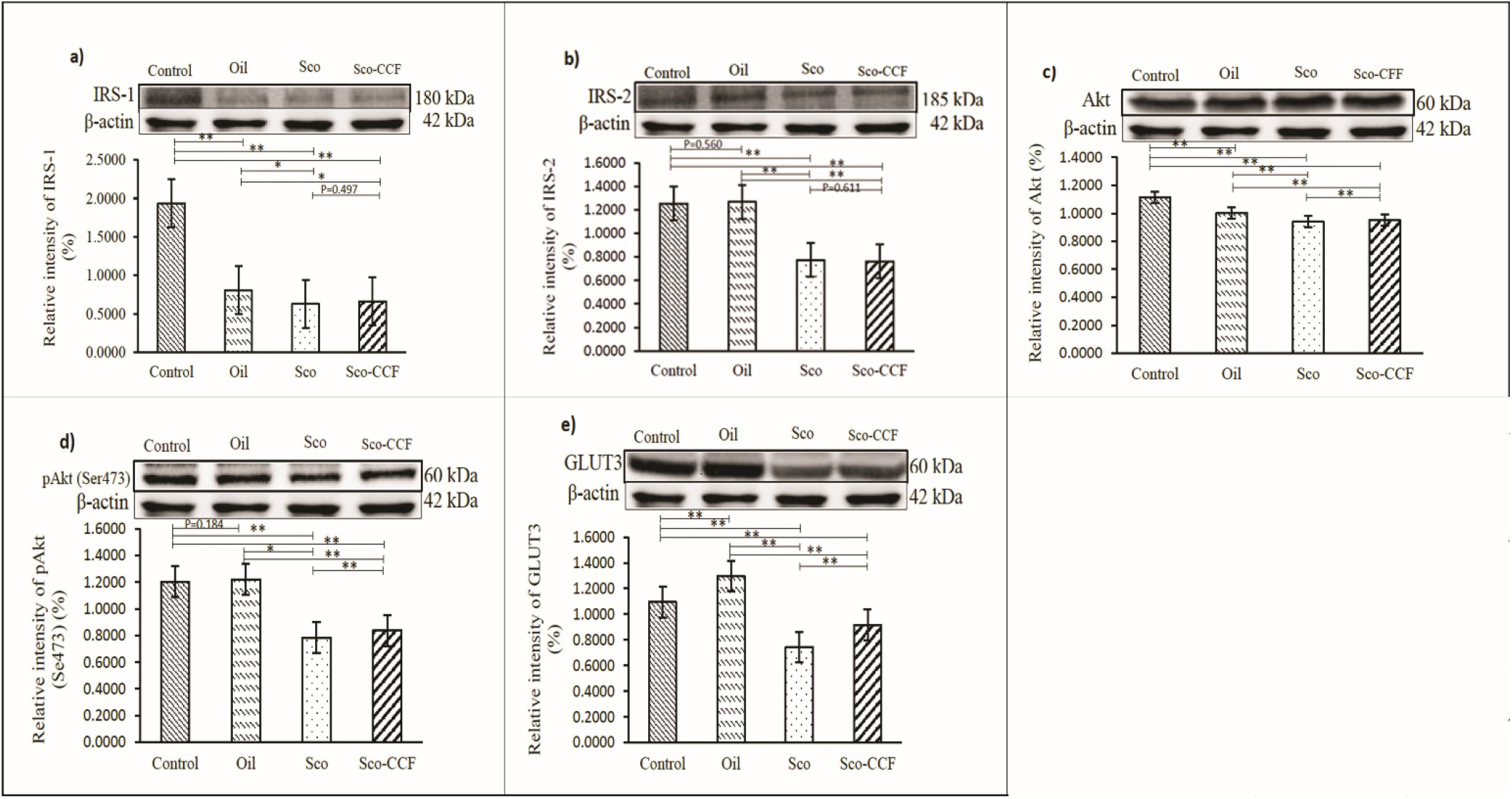
Immunoblotting and densitometry analysis of (a) IRS-1 (b) IRS-2 (c) Akt (d) pAkt (Ser 473) and (e) GLUT3 in the brains (n=3) in brain lysate. The relative fold of change in the levels of proteins were normalized with respect to the level of β-actin, which was used as a loading control. Data expressed as mean ± SD. Control: saline-treated, Oil: Sunflower oil treated, Sco: Scopolamine and Sco-CCF: Scopolamine-cholecalciferol treated. *P < 0.01 and **P < 0.001.

### Reduced levels of glucose transporter GLUT3 in AD rat brain and cholecalciferol significantly reversed it

We investigated the expression of a major glucose transporter GLUT3 among the various groups in the rat brains quantitatively using immunoblotting (Figure 3e). In comparison to saline-treated control group, the expression of GLUT3 was significantly reduced (P<0.001) in scopolamine-induced AD brain (Figure 2). However, cholecalciferol significantly increased GLUT3 (P<0.001) expression in AD rat brain. Interestingly, expression of GLUT3 was significantly different (P<0.001) between saline-treated control group and sunflower oil-treated group (Figure 3e).

## Discussions

Insulin signaling pathway is an important biochemical pathway at the cellular level and affects cellular homeostasis. From previous studies, it was found that lower expressions of IRs, IGF-1Rs, and IRS expression were associated with AD. The decrease of IRS-1/2 levels indicated that insulin transportation into the brain was impaired [22,25]. Other studies suggested that declined IRS-1 expression may be associated with reduced memory performance in rats [26]. Here, we observed that the levels of upstream insulin signaling molecules in the IRS-1 and IRS-2 pathways were markedly decreased in the scopolamine-induced AD rat model, compared to the saline-treated-control rat brain (Figure 3).

Akt also has multiple phosphorylation sites generally, phosphorylation at Ser473 requires the full activation of Akt and so, the level of phosphorylation at Ser473 represents the degree of Akt phosphorylation [27]. Previous studies suggested that the activation of Akt can prevent neuronal loss in AD and it was reported that there was a reduced localization of cells immunostained for pAkt (Ser473) in AD brains [22,28]. In our study, in scopolamine-induced rats, we observed that there was a reduction in the localized of cells immunostained for Akt and pAkt (Ser473) compared to the saline-treated control rat (Figure 2). We observed a significant decline in the expression of pAkt (Ser473) and Akt in scopolamine-induced rats, in comparison to saline-treated control rats (Figure 3). In addition, another study has investigated the association of Akt expression and memory performance in scopolamine-induced animals and they reported that there was a decline in Akt expression in the brain in scopolamine-induced animals that exhibited memory impairment [29]. Akt was believed to be an important contributor to scopolamine-induced memory impairment. This means that our study in scopolamine-induced rats, showing reduced amount of Akt and pAkt (Ser473) may be associated with a reduced spatial memory performance (Figures 1).

It was well known that GLUT3 is the main neuronal glucose receptor in the brain and it has a higher glucose affinity and more potential than GLUT1 for glucose transportation [30]. Therefore, it is believed that higher GLUT3 expression might improve glucose uptake in the neuron [31]. From previous studies, it was found that the level of GLUT3 was decreased in the AD-affected brain; suggesting decreased GLUT3 may contribute to AD neurodegeneration. In our study, in scopolamine-induced rats, we observed the immunostained cells for GLUT3 were rescued, compared to saline-treated-rat (Figure 2). In addition, insulin binding triggers tyrosine phosphorylation of the insulin receptor and IRSs leading to the activation of PI3K, Akt, and pAkt (Ser473) which promote expression of GLUT3 for glucose uptake and induce insulin release from the cells [30,31]. Disruption in the IRSs and Akt activation and, subsequent cascade of GLUT3 may result in impaired insulin signaling, thus indirectly resulting in abnormal elevation of insulin which promotes insulin resistance, associated with AD. It was suggested that reduced Akt-mediated activation of GLUTs and declined GLUTs expression resulted in impaired insulin signaling cascade in the AD brain [31]. Impaired insulin signaling leading to impaired glucose metabolism in AD was suggested due to decreased expression of GLUT3. Other studies showed that the interaction of IRSs with PI3 kinase was necessary for the activation of GLUT3 in the brain. From the densitometry analysis, we observed a significant decline in the expression of GLUT3 in the brains of scopolamine-induced rats, in comparison to the saline-treated rat (Figure 3).

Therefore, our study showed, in comparison to the saline-treated rat brain, a declined memory performance in scopolamine-induced which may be due to reduced expression of IRS-1 and IRS-2, p-Akt (Ser473), Akt, which are highly associated with reduced GLUT3 expression. This might be due to the preceding reduction in the IRS-1, IRS-2, PI3K, Akt pathways leading to impairment in the insulin signaling and one that might promote insulin resistance associated with AD, which might create the hypoglycemia in the brain by the dysfunctional of PI3k dependent pathway via Akt (Figure 4).

**Figure 4.**
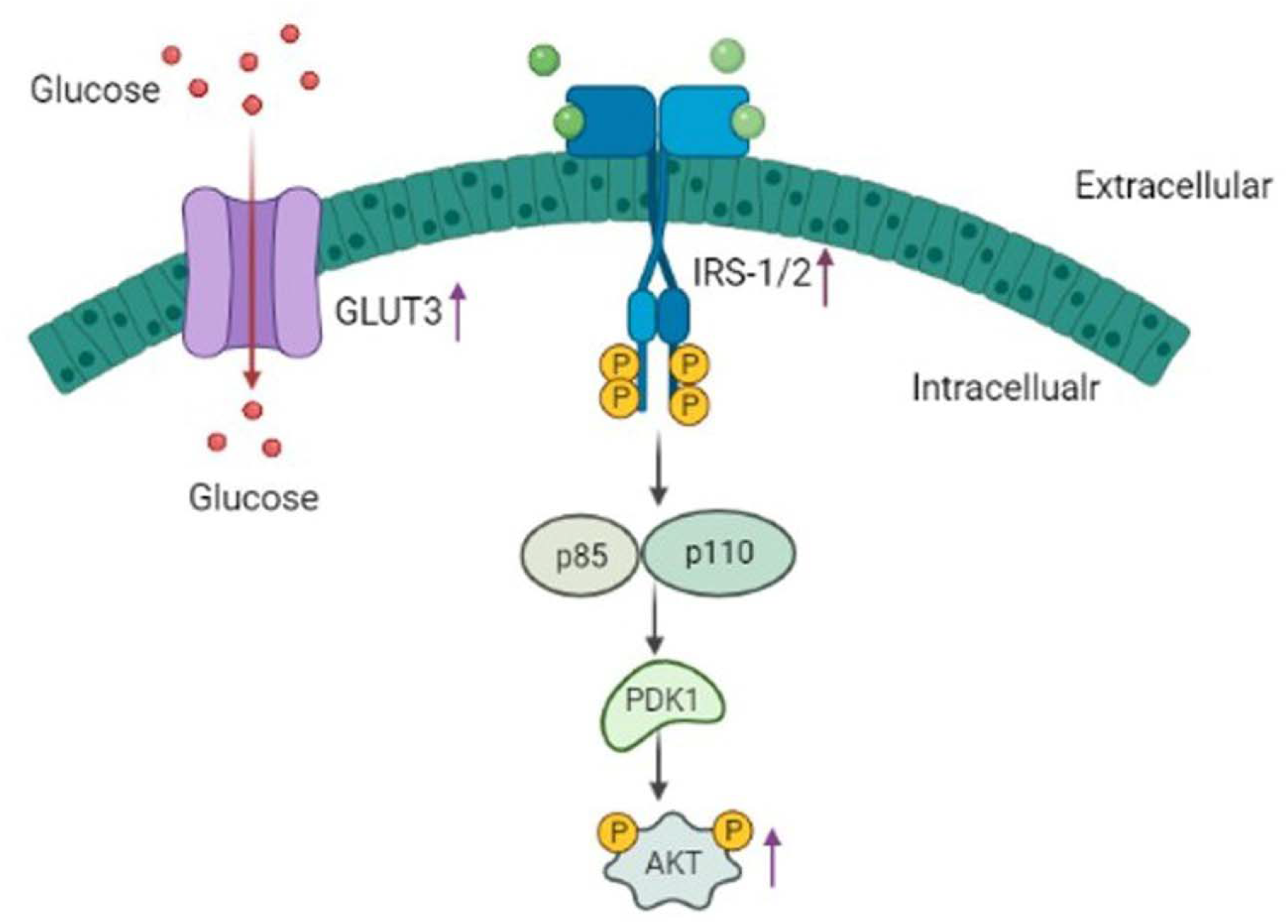
Schematic representation for the mechanism of action of cholecalciferol in treatment of AD. Cholecalciferol increases IRS1/2, Akt, and p-Akt (Ser 473) in AD rats causing reduced aggregation of Amyloid beta in the brain. In addition, expression of GLUT3 was increased (purple arrow) by treatment of cholecalciferol in AD rats.

In addition, here Sunflower oil was used as a solvent for cholecalciferol. Although, Sunflower oil is one of the most widely used vegetable oil for human consumption by volume after palm and soybean oil worldwide. However, limited scientific evidences would suggest some benefits for olive oil consumption in different mouse models of neurodegeneration [32, 33]. By contrast, no data are available on the effect that sunflower oil consumption on AD pathogenesis. For this reason, in this research work we assessed the biological effect of sunflower oil administration on scopolamine-induced AD rats. Based on our study, sunflower oil may have some beneficial effects. It may be associated with the amelioration of insulin signaling via mediating upregulation of IRS-1, Akt, and GLUT3 expression in the brain. Further studies are warranted to better understand the mechanism of action of sunflower oil in the treatment of AD.

## Conclusion

In summary, reduced expression of IRS-1 and IRS-2 in scopolamine-induced rats can lead to reduced phosphorylation of Akt and pAkt (Ser473) which are associated with reduced GLUT3 expression which declined spatial memory performance which all are associated with the development of AD. Our study suggests that cholecalciferol may ameliorate impaired insulin signaling via upregulation of IRS-1, IRS-2, Akt, pAkt (Ser473), and GLUT3 expression. Thus, cholecalciferol treatment can an alternative therapeutic treatment for preventing and/or delaying the development of AD.

## Author Contributions

T.K.D. and M.W.R. involved in the study design; T.K.D. and E.Y. completed the experiment; T.K.D. and E.Y. performed data analysis and interpretation; T.K.D. and E.Y. wrote the original draft.; T.K.D. and M.W.R edited the manuscript. All authors have read and agreed to the published version of the manuscript.

## References

1. Alzheimer’s Association Report. 2021 Alzheimer’s disease facts and figures. Alzheimers Dement. 2021, 17 (3), 327–406.

2. Alzheimer’s Association Report. 2014 Alzheimer’s disease facts and figures, Alzheimers Dement. 2014, 10 (2), e47–e92.

3. Sato Y, Asoh T, Oizumi K. High prevalence of vitamin D deficiency and reduced bone mass in elderly women with Alzheimer’s disease. Bone. 1998; 23(6), 555–557.

4. Khanh VQL, Lan THN. The Beneficial Role of Vitamin D in Alzheimer’s Disease. American Journal of Alzheimer’s Disease & Other Dementias. 2011, 26 (7), 511–520.

5. Lukaszyk E, Bien-Barkowska K, Bien B. Cognitive functioning of geriatric patients: is hypovitaminosis D the next marker of cognitive dysfunction and dementia? Nutrients. 2018, 10(8), 1104.

6. Sadia S, Uzma T, Shatha AB, Waad KAO, Afaf A, Fatimah A, Amira H. Low Vitamin D and Its Association with Cognitive Impairment and Dementia. Journal of Aging Research. 2020,1–10.

7. Duygu GA, Erdinç D, Turan E, Haşmet H, Hakan G, Murat E, Engin E, Melek O, Funda E, Selma Y. Association between vitamin D receptor gene polymorphism and Alzheimer’s disease. Tohoku J Exp Med. 2007, 212(3),275–82.

8. Nanyang L, Tingting Z, Lina M, Wei W, Zehui L, Xuefan J, Jiahui S, Hui P, Hao L. Vitamin D Receptor Gene Polymorphisms and Risk of Alzheimer Disease and Mild Cognitive Impairment: A Systematic Review and Meta-Analysis, Advances in Nutrition, 2021.

9. Darryl WE. Vitamin D: Brain and Behavior. JBMR. 2021, 5 (1).

10. Nathanael J Y, Dijana T, Kirk WF, Jeremy TS, Michael WC, Celeste W, Rachael CC, Michaela DW, Andrew JOW, Caitlin SW. Vitamin D is crucial for maternal care and offspring social behaviour in rats. Journal of Endocrinology. 2018, 237, 73–85s.

11. Milou AP, Elske MBB. The Impact of Maternal Vitamin D Status on Offspring Brain Development and Function: A Systematic Review. Adv Nutr. 2016,7(4),665–678.

12. Wei L, Luke AH, Stephanie V, Urs M, Darryl WE. Maternal Vitamin D Prevents Abnormal Dopaminergic Development and Function in a Mouse Model of Prenatal Immune Activation. Sientific Reports, 2018,8:9741,1-12.

13. Ricardo GO, Noelia GD, Samuel DG, Livia C, Cristina V, Pedro NA, Carmen C. Vitamin D deficiency as a potential risk factor for accelerated aging, impaired hippocampal neurogenesis and cognitive decline: a role for Wnt/β-catenin signaling. Aging. 2020, 12 (13). 13824–13844.

14. Maria M, Véréna L, Emmanuelle L, Kevin B, Cedric A, François F, Pascal M. Vitamin D Improves Neurogenesis and Cognition in a Mouse Model of Alzheimer’s Disease. Molecular Neurobiology, 2018, 55:6463–6479.

15. Hypponen E, Laara E, Jarvelin MR, Virtanen SM. Intake of vitamin D and risk of type 1 diabetes: a birth-cohort study. Lancet, 2001, 358, 1500–1503.

16. Mitri J, Muraru MD, Pittas AG. Vitamin D and type 2 diabetes: a systematic review. European Journal of Clinical Nutrition, 2011, 65, 1005–1015.

17. Shamaila R, Per B J. Is Hypovitaminosis D Related to Incidence of Type 2 Diabetes and High Fasting Glucose Level in Heal 2018, 10(1), 59.1-18.

18. Izabela SP, Agnieszka S. Analysis of Association between Vitamin D Deficiency and Insulin Resistance. Nutrients. 2019, 11, 794, 1–28.

19. Christian T, Verena S, Marlene P, et al.: Effects of Vitamin D Supplementation on IGF-1 and Calcitriol: A Randomized-Controlled Trial. Nutrients. 2017, 9(6), 623.1-10.

20. Das T K, Chakrabartiby SK, Zulkiplia IN and Mas RW. Curcumin Ameliorates the Impaired Insulin Signaling Involved in the Pathogenesis of Alzheimer’s Disease in Rats. Journal of Alzheimer’s Disease Reports, 2019, 3,59–70.

21. Sami G, Simo R, Mikael M, et al. Altered Insulin Signaling in Alzheimer’s Disease Brain – Special Emphasis on PI3K-Akt Pathway. Front. Neurosci. 2019,18.1–8.

22. Das TK, Jana P, Chakrabarty SK Mas RW. Curcumin Ameliorates the Impaired Insulin Signaling Involved in the Pathogenesis of Alzheimer’s Disease in Rats. Journal of Alzheimer’s Disease Reports. 2019, 3,59–70.

23. Haider S, Tabassum S, Perveen T. (2016) Scopolamine induced greater alterations in neurochemical profile and increased oxidative stress demonstrated a better model of dementia: A comparative study. Brain Res Bull. 2016, 127, 234–247.

24. Dasgupta, S., Rai, R.C. PPAR-γ and Akt regulate GLUT1 and GLUT3 surface localization during Mycobacterium tuberculosis infection. Mol Cell Biochem 440, 127–138 (2018).

25. Matteo S, Salvatore F and Claudio G: Brain Insulin Resistance and Hippocampal Plasticity: Mechanisms and Biomarkers of Cognitive Decline. Front. Neurosci. 2019:1–13.

26. Hilaree NF, Katie LA, Adam OG, et al.: Molecular elevation of insulin receptor signaling improves memory recall in aged Fischer 344 rats. Aging Cell. 2020;19:e13220.1-13.

27. Muheeb B, Nazish A, Fathima ST, Nasser KA, Timothy EM. Distinct Akt phosphorylation states are required for insulin regulated Glut4 and Glut1-mediated glucose uptake. Cite this articleas: eLife 2017;6:e26896 DOI: 10.7554/eLife.26896

28. Sami G, Simo R, Mikael M, et al.: Altered Insulin Signaling in Alzheimer’s Disease Brain – Special Emphasis on PI3K-Akt Pathway. Frontiers in Neuroscience. 2019.13:1–8.

29. Yulu W, Shichao G et al.: A Novel PDE4D Inhibitor BPN14770 Reverses Scopolamine-Induced Cognitive Deficits via cAMP/SIRT1/Akt/Bcl-2 Pathway. Front. Cell Dev. Biol., 10 December 2020:1–13.

30. Hermann K. Glucose transporters in brain in health and disease. Pflügers Archiv - European Journal of Physiology (2020) 472:1299–1343.

31. Natalia K, Hedley CAE, Natalia K, Hedley CAE, et al.: A Systematic Review of Glucose Transport Alterations in Alzheimer’s Disease. Front. Neurosci., 20 May 2021:1–15.

32. Farr S A et al. Extra virgin olive oil improves learning and memory in SAMP8 mice. J Alzheiermer’s Dis. 2012, 28, 81–92.

33. Pitozzi, V et al. Effects of dietary extra virgin olive oil on behavior and brain biochemical parameters in ageing rats. Br J Nutr. 2010, 103, 1674–1683.

